# Sequences in the Human papillomavirus Type 18 (HPV18) Upstream Regulatory Region regulate Viral Genome Replication, Establishment and Persistence

**DOI:** 10.1101/2021.02.22.432334

**Authors:** Tami L. Coursey, Koenraad Van Doorslaer, Alison A. McBride

## Abstract

During persistent human papillomavirus infection, the viral genome replicates as an extrachromosomal plasmid that is efficiently partitioned to daughter cells during cell division. We have previously shown that an element which overlaps the HPV18 transcriptional enhancer promotes stable DNA replication of replicons containing the viral replication origin. Here we perform comprehensive analyses to elucidate the function of this maintenance element. We conclude that no unique element or binding site in this region is absolutely required for persistent replication and partitioning, and instead propose that the overall chromatin architecture of this region is important to promote efficient use of the replication origin. These results have important implications on the genome partitioning mechanism of papillomaviruses.

## Introduction

High-risk, oncogenic human papillomaviruses (HPVs) represent a global health burden; they are the etiological agents of several cancers, causing ~5% of all human cancers [1]. The oncogenic potential of these high-risk HPVs is tied to their lifecycle in keratinocytes. HPV infects proliferative, basal layer keratinocytes that support viral genome replication [2]. Extrachromosomal HPV genomes (minichromosomes) replicate at a low level in these dividing cells and are thought to be partitioned into daughter cells by tethering to host mitotic chromosomes [3]. The high-risk E6/E7 oncogenes interfere with many cellular processes to produce an environment conducive for persistent viral infection. However, longterm abrogation of cell cycle checkpoints during persistent infection can promote oncogenesis [4]. Thus, defining the mechanism behind HPV genome maintenance could provide insight into therapeutics to treat persistent HPV infection.

Viral genome replication and partitioning of papillomaviruses requires the E1 and E2 replication proteins, the origin of replication, and additional sequences from the Upstream Regulatory Region (URR) that promote long-term genome replication and partitioning [5–7]. The origin of replication contains an E1 binding site, flanked by E2 binding sites (E2BSs) and cooperative binding of the E1 helicase and E2 proteins to these sites drives initiation of viral DNA synthesis, but is insufficient for long-term maintenance of the viral minichromosomes. In bovine papillomavirus 1 (BPV1), genome maintenance requires both the origin and a region from the URR that overlaps the enhancer and was designated the Minichromosome Maintenance Element (MME) [7]. In BPV1, the MME encompasses approximately six E2 binding sites upstream from the origin (for a total of eight E2BSs) [7]. However, the URRs of Alphapapillomavirus HPVs contain only four E2BSs, and we have shown previously that the E2BS most distal from the origin is not required for genome maintenance [8]. Instead, we showed that for HPV18, a core element from the enhancer region of the URR (nt 7564-7641), in addition to the origin of replication, was required for genome maintenance in proliferating keratinocytes [8], and we designated this the MMEE; minichromosome maintenance enhancer element [3]. Additionally, Ustav and colleagues have shown that only two E2BSs are required for persistence of HPV18 DNA in the absence of DNA replication [9]. Therefore, the partitioning requirements for the oncogenic HPVs are still unclear.

In this study, we further explored the requirement and function of the core enhancer region (MMEE) in HPV18 replicon maintenance. We use two stringent assays that measure the ability of replicons to replicate, establish as extrachromosomal plasmids, and persist in primary keratinocytes. In the first, the replicon is transfected into primary keratinocytes along with the HPV18 genome and the cells are passed five times in the absence of any selection. The HPV18 genome expresses the E1 and E2 replication proteins, while the E6 and E7 proteins additionally provide a selective growth advantage to the transfected cells. Only replicons that can replicate and partition successfully are retained alongside the HPV18 genome over multiple cell passes. Viral DNA is measured by qPCR and Southern blot analysis; these assays provide a precise measure of replicon stability and copy number over time. In the second assay, the co-transfected cells are grown under G418 selection and those cells in which the replicon replicates, and persists, grow to form a drug-resistant colony. This provides a quantitative measurement of the efficiency by which the replicon can establish in a cell while the growth and size of the colony is a readout of persistence and partitioning. This assay does not give much information about replicon copy number, but replicons that persist at very low copy number do give rise to stable G418 resistant colonies. Lastly, the pCGneo replicon can replicate and express the drug resistance marker in both prokaryotic and eukaryotic cells; thus, it can be isolated from keratinocytes and transformed into bacteria for further evaluation.

In this study, we use these assays to further evaluate and identify sequences from the HPV18 core enhancer that are critical for persistent replication of HPV18 derived replicons. We conclude that there is no single element within this region that is absolutely essential for persistence but rather that these sequences collectively promote persistent replication by facilitating the formation of beneficial chromatin structure around the origin.

## Results

### Delineation of Sequence Elements in the HPV18 Upstream Regulatory Region

To explore the elements in the HPV18 URR that support and modulate maintenance replication, we used the complementation assay that we previously developed in which primary human foreskin keratinocytes (HFKs) are co-electroporated with the HPV18 genome and a replicon containing both the origin of replication and elements from the URR required to promote long-term replication [8]. The pCGneo replicon is a CpG-free plasmid with minimal bacterial elements and a neomycin resistance cassette that is expressed in both bacterial and eukaryotic cells (Figure 1A). Replicon maintenance over long-term cell division can be assessed in the presence and absence of G418 selection. In our previous study, we showed that part of the core enhancer from the central URR was required for maintenance of the replicon in addition to the replication origin (Figure 1B) [8]. This 77bp region (nt 7568-7642), designated Region 2 or the MMEE, was indispensable for long-term replicon maintenance, while the flanking Regions 1 and 3 were not required. Region 2 contains binding sites for Nuclear factor-1 (NF-1), Yin-yang 1 (YY1) and Activator protein 1 (AP1) [10, 11] that are essential for regulation of the P105 early promoter [12, 13].

**Figure 1:**
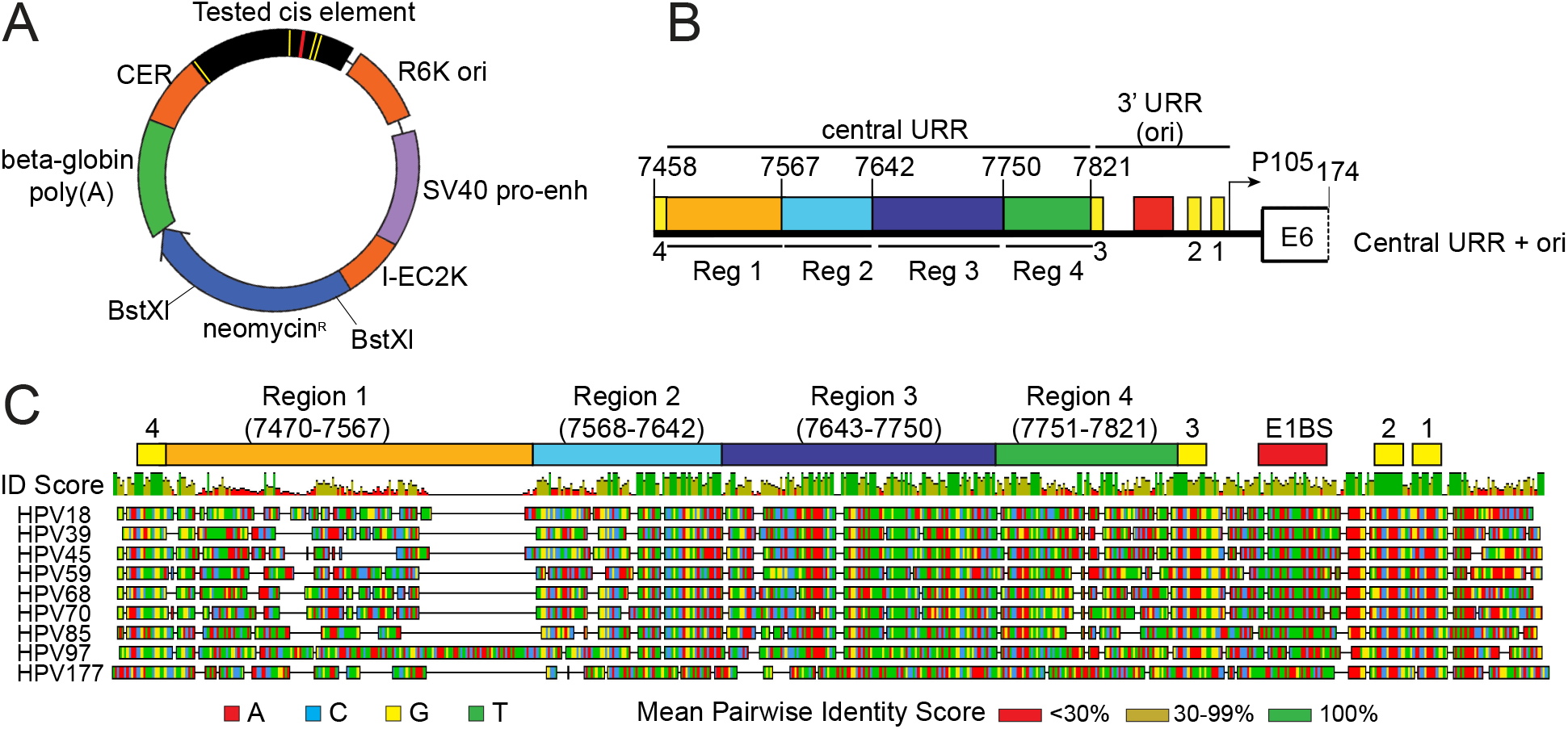
Diagram of the HPV18 replicon and the HPV18 URR cis element insertion. A. Diagram of the HPV18 replicon plasmid, pCGneo. Prokaryotic elements are highlighted in orange (R6K ori, I-EC2K, and CER) while eukaryotic elements are highlighted in violet (simian virus 40 [SV40] promoter-enhancer) and green (beta-globin polyA signal). The neomycin resistance gene is in blue. The HPV18 Central URR and origin are represented in black with E1 and E2 binding sites indicated by red and yellow bars, respectively. The *BstX*l restriction enzyme sites used for Southern blot analysis are indicated. B. Diagram of the HPV18 Central URR and 3’URR origin sequences inserted into the replicon. The four regions of the Central URR are indicated, and nucleotide numbers are from the HPV18 genome. The arrow represents the early viral P105 promoter. C. A global alignment of the central URR and ori from the nine Alphapapillomavirus 7 species of human papillomaviruses using the Geneious Prime multiple sequence alignment algorithm of the central and 3’-terminal half of the URR. The HPV18 central URR and origin elements are indicated. Nucleotides within the sequences are color coded as indicated. A mean pairwise identify score (sliding window size of 1) for each nucleotide is shown graphically above the aligned sequences and indicate less than 30% (red), between 30 to 99% (yellow), or 100% nucleotide conservation among the sequences.

To further investigate sequences in the URR involved in replicon maintenance, we aligned the HPV18 central URR sequences with those from eight other Alphapapillomavirus 7 HPV species. This alignment showed high sequence identity in the 3’ half of Region 2, Region 3, and also in an additional 67bp region between Region 3 and E2BS3 in the origin (Region 4, nt 7751-7821) (Figure 1C). We had previously shown that Region 4, in combination with the minimal origin, is unable to support replicon maintenance [8], but it remained unclear whether it contributed to replication maintenance. Notably, Region 4 contains a binding site for AP1 which is necessary to promote transcription from the HPV18 early viral P105 promoter [13].

### Region 4 of the HPV18 URR, adjacent to the minimal replication origin, is not required for replicon maintenance

To explore the requirement of Region 4 in replicon maintenance, a series of pCGneo nt 7452-174 Central URR+ori replicons were generated with partial (nt 7751-7764 or 7772-7821) or full (nt 7750-7821) deletions of Region 4 (Figure 2A). Keratinocytes were co-electroporated with the replicons and the HPV18 genome and selected with G418 until colonies formed. Keratinocytes transfected with all replicons formed robust neomycin resistant colonies, comparable to the pCGneo 7452-174 (Central URR+ori) replicon. Therefore, Region 4 is unnecessary for HPV18 replicon establishment and maintenance (Figure 2B).

**Figure 2.**
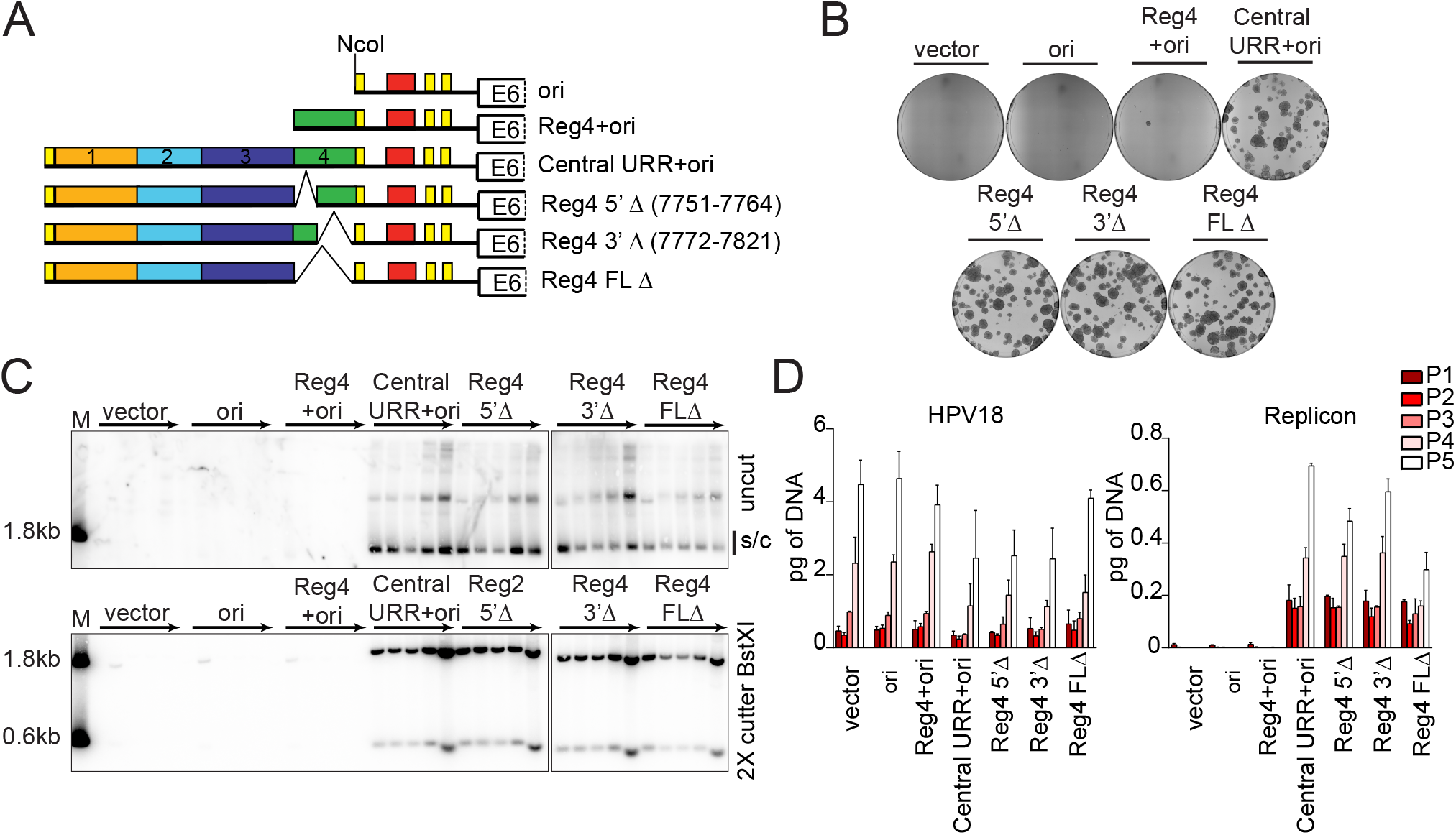
HPV18 central URR element, Region 4, is unnecessary for replicon maintenance. A. Diagram of replicons containing the Central URR and origin with deletions in Region 4 (green). Control replicons include the CpGneo (vector), the CpGneo vectors with the minimum HPV18 origin (ori), Region 4 and the origin (Reg4+ori), and the central and origin (Central URR+ori). B. Keratinocyte colonies arising from continuous G418 selection were stained with methylene blue approximately 14 days post-transfection. C. Cellular DNA collected from five passages of co-transfected cells were digested and analyzed by Southern blot and viral DNA detected by hybridization with a radioactively labeled CpGneo vector probe. The top panel represents DNA samples cut with an enzyme that does not cut the replicon, permitting visualization of extrachromosomal, supercoiled replicon DNA. The bottom panel shows DNA cut with *BstX*l which separates the replicon into two fragments. Lanes containing a size marker (M) contain two fragments generated from digesting the CpGneo vector with *BstX*l. An asterisk marks cellular DNA hybridizing with the beta-globin polyA region present in the radioactive CpGneo vector probe. The monomeric, supercoiled form (s/c) of the replicon is indicated. D. HPV18 genome (left) and replicon (right) copy numbers were measured by qPCR. Data shown is representative (B, C), or an average of two biological replicates (D).

To ensure that the replicons were maintained extrachromosomally during long-term cell division, and to measure replicon copy number, the co-transfected keratinocytes were cultured without selection for five passes (~15-18 population doublings). Total DNA extracted from each pass was analyzed by Southern blot for replicon maintenance (Figure 2C). Deletion of any part of Region 4 did not impact extrachromosomal maintenance of the replicon, although replicons with the full deletion (nt 7752-7821) had slightly reduced copy number.

HPV18 genome and replicon copy numbers were also measured by quantitative PCR (qPCR) (Figure 2D) in the DNA samples from long-term passage. The HPV18 copy number was relatively consistent in all co-transfections and increased per pass. This increase is due to immortalization of the primary keratinocytes by HPV18, which gives a selective growth advantage to transfected cells. Concomitantly, the copy number of the HPV18 replicons also increased per pass except in the case of the full-length Region 4 deletion. However, the levels of this replicon were stable from pass three to five. Thus, although Region 4 is unnecessary for replicon maintenance it may play a role in bolstering the copy number and/or partitioning efficiency for maintenance during long-term cell division.

### HPV18 URR Region 2 is necessary, but not sufficient, to support maintenance of replicons containing the minimal replication origin

The data above support previous findings that Region 2 is the only URR element necessary for replicon maintenance [8]. Nevertheless, other URR elements may contribute to replicon maintenance by regulating copy number and/or partitioning efficiency. To determine whether Region 2 is sufficient for replicon maintenance, Region 2 was inserted upstream from the minimal origin either with or without Region 4 (replicons Reg2+4+ori and Reg2+ori, respectively) (Figure 3A). As shown in Figure 2, Region 4 was neither sufficient, nor required, to support maintenance of replicons containing the minimal replication origin. Keratinocytes were co-transfected with HPV18 DNA and the pCGneo replicons and grown under G418 selection. The Reg2+4+ori replicon was maintained and supported robust, expanding neomycin resistant keratinocyte colonies. In contrast, keratinocytes co-transfected with Reg2+ori replicons formed only abortive/collapsing colonies (Figure 3B) indicating while the replicon could replicate initially, and express the neomycin resistance gene, it was not stably partitioned into daughter cells.

**Figure 3.**
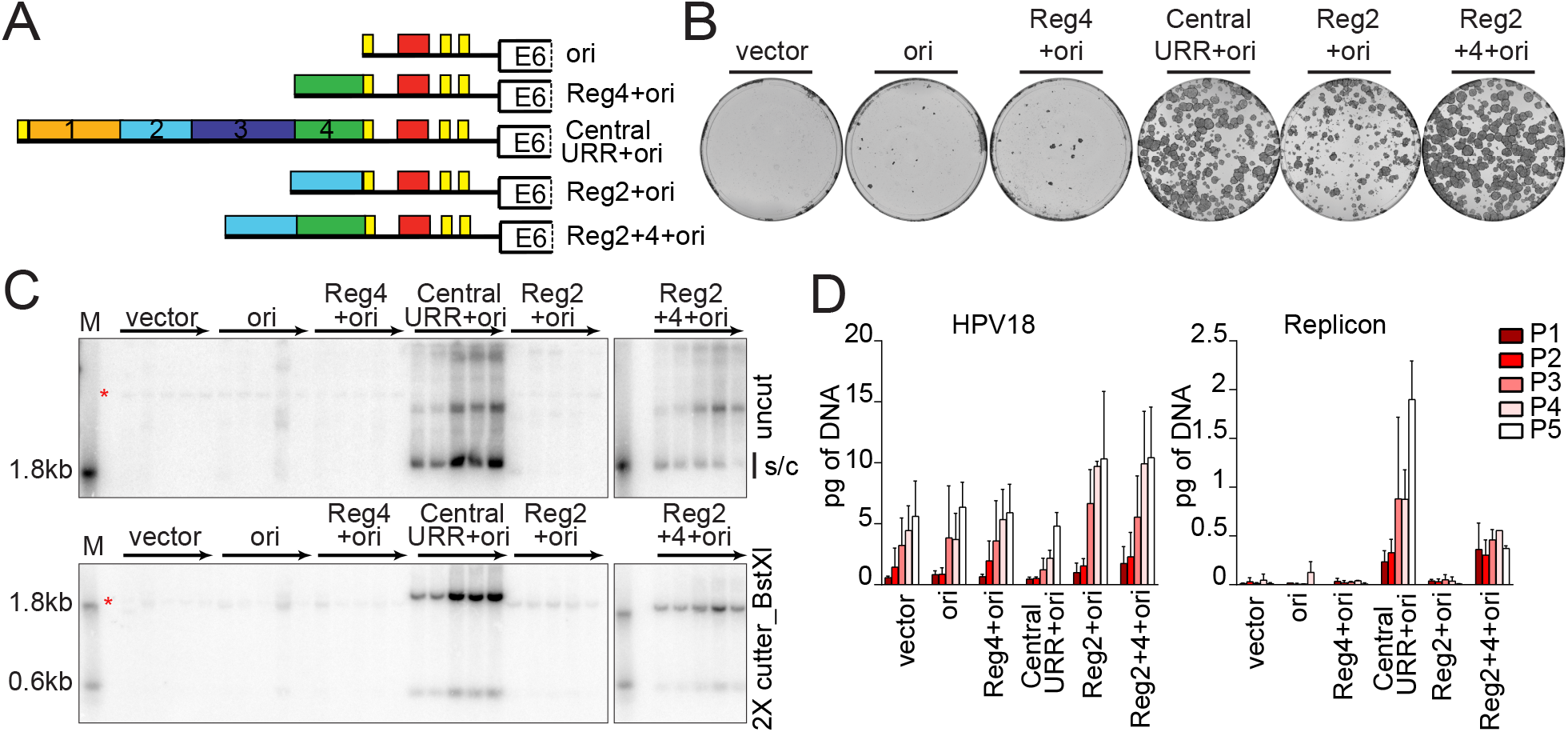
HPV18 central URR element Reg2 alone is necessary but insufficient for replicon maintenance. A. Diagram of HPV18 replicons. B. Keratinocyte colonies arising from continuous G418 selection were stained with methylene blue approximately 14 days post-transfection. C. DNA collected from five passages of transfected cells were analyzed by Southern blot as described in Figure 2C. An asterisk marks cellular DNA hybridizing with the beta-globin polyA region present in the radioactive CpGneo vector probe. The monomeric, supercoiled form (s/c) of the replicon is indicated. D. HPV18 genome (left) and replicon (right) copy numbers were measured by qPCR. Error bars represent standard deviation. Data shown is representative (B, C), or an average of two biological replicates (D).

To confirm this defect in replication, and to determine whether manipulation of the HPV18 enhancer element might affect expression of the neomycin resistance gene, transfected keratinocytes were also cultured for five passes in the absence of G418 selection and extrachromosomal maintenance of the replicon was assessed by Southern blot analysis (Figure 3C). In support of the colony formation results (Figure 3B) only replicons containing the central URR and Reg2+4+ori were efficiently maintained extrachromosomally for five passages. qPCR measurements of both HPV18 and replicon copy number supported the Southern blot analysis (Figure 3D). However, while the Reg2+4+ori replicon was maintained extrachromosomally, its overall copy number did not increase with pass, and over time the monomer form was reduced with a concomitant increase in multimeric forms (Figure 3C). Thus, additional elements from the central URR (e.g. Reg1 or 3) may contribute to more robust maintenance by increasing replicon copy number and/or partitioning efficiency. Moreover, because our results in Figure 2 demonstrated that sequences in Region 4 were unnecessary for replicon maintenance, the defect in maintenance of the Reg2+ori replicon could indicate that the nucleotide spacing between Region 2 and the origin of replicon is critical. The close spacing of the origin and enhancer regions could cause steric hinderance of binding of cellular and/or viral factors to these regions.

### Enhancer elements cannot substitute for Region 2 in replicon maintenance

In SV40, transcriptional enhancer elements augment the efficiency of viral DNA replication [14, 15]. Region 2 is located in the core of the HPV18 enhancer and contains multiple binding sites for factors important for transcriptional regulation [16]. Therefore, the role of Region 2 in replicon maintenance could be to recruit cellular transcription factors and/or alter the chromatin structure of the viral minichromosome. To determine whether enhancer activity per se supports replicon maintenance, three different enhancers were inserted upstream from Reg4+ori (Figure 4A). These enhancer elements were introduced upstream of Region 4 to reduce possible steric hindrance between the enhancers and the origin region. Two of these elements are cellular enhancers that drive ubiquitous expression of the human aldolase A gene (hAldA) or keratinocyte specific expression of the keratin 5 (K5) gene [17–19]. The K5 enhancer was cloned in forward or reverse orientations (designated as K5^enh^-F or -R) in the replicon to control for any unknown orientation specific impacts on enhancer activity. The third enhancer, from the human cytomegalovirus (HCMV), promotes strong and ubiquitous activity. These enhancers exhibit activity in diverse cell types and contain transcription factor binding sites that are also present in the HPV18 central URR (e.g. AP1) [17, 19, 20].

**Figure 4.**
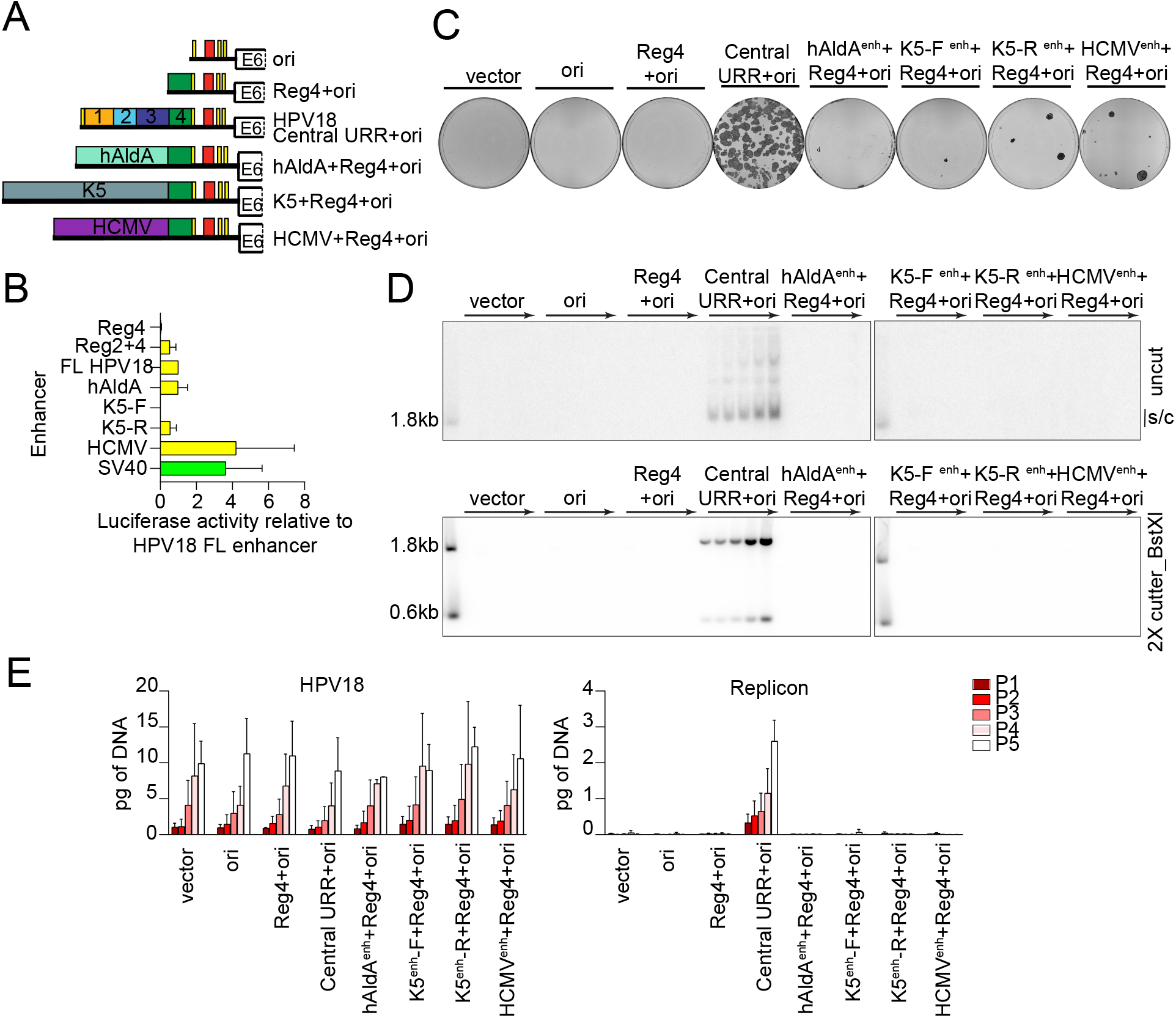
Enhancer activity alone cannot complement the HPV18 central URR enhancer element for replicon maintenance. A. Diagram of enhancer replicons. B. Keratinocytes were transfected with luciferase reporter plasmids containing the cis elements indicated upstream from a minimal promoter, as well as a positive control (pGL4.13 expressing luciferase from the SV40 enhancer/promoter; shown in green) and negative control (empty luciferase vector containing a minimal promoter). Cell lysates were collected 48 hours after transfection and luciferase activity was measured and normalized to the total protein concentrations for each cell lysate and the negative control plasmid. Reporter activity for each enhancer is shown relative to the HPV18 central URR enhancer. C. Keratinocyte colonies arising from continuous G418 selection were stained with methylene blue approximately 14 days post-transfection D. DNA collected from five passages of co-transfected cells were analyzed by Southern blot as described in Figure 2C. The monomeric, supercoiled form (s/c) of the replicon is indicated. E. HPV18 genome (left) and replicon (right) copy numbers were measured by qPCR. Error bars represent standard deviation. Data shown are representative (C, D) or an average of two (E) biological replicates. The average of three biological replicates is shown in B.

To confirm the enhancer activity in keratinocytes, each element was also cloned upstream from a minimal promoter in a luciferase reporter (Figure 4B). The HPV18 central URR enhanced luciferase expression ~ 41-fold compared to the minimal promoter. In comparison, Region 2 and 4, while sufficient for replicon maintenance (Figure 3), enhanced luciferase expression only ~17 fold. The hAldA enhancer had activity comparable to the HPV18 central URR (~33 fold), while the K5 enhancers enhanced luciferase expression 0.5 to 18.4-fold, depending on orientation. In comparison, the SV40 enhancer (in a positive control plasmid) and HCMV enhancer activity enhanced luciferase expression 119 and 129-fold, respectively. Despite their range of enhancer activity, the pCGneo enhancer replicons failed to give rise to robust neomycin resistant colonies when co-transfected into keratinocytes with the HPV18 genome. (Figure 4C). Therefore, enhancer strength does not correlate with the ability to support colony formation (at least in the absence of replicon maintenance).

Southern blot analysis of DNA isolated from keratinocytes transfected with the test replicons (and HPV18) and passaged in the absence of G418 selection, showed, despite the range of enhancer activities, none of the test replicons were maintained extrachromosomally during long-term cell division (Figure 4D). Measurement of the co-transfected HPV18 genome and replicon DNA copy numbers confirmed that while the HPV18 genome is maintained at similar copy numbers among samples, the test replicons are not (Figure 4E). Therefore, strong enhancer activity cannot substitute for the replicon maintenance function of Region 2 or the central URR; thus, the mechanism of replicon maintenance is not exclusively due to the enhancer activity of the HPV18 central URR elements.

### HPV16 and HPV31 Central URR regions minimally support HPV18 replicon maintenance

The results above suggested that there was no simple correlation between enhancer activity and the ability to support replicon maintenance. Instead, the enhancer region of HPV18 must support replicon maintenance by a more complex mechanism. To determine whether the homologous enhancers from other HPVs could support persistent, extrachromosomal maintenance, we substituted the E2BS4 and central URR regions (upstream of E2BS3) from Alphapapillomavirus 9 species HPV16 (nt 7451-7867) and HPV31 (nt 7477-7867) into the equivalent position in the HPV18 URR in pCGneo (HPV18 nt 7458-7821) (Figure 5A). Extrachromosomal maintenance was assessed by co-transfecting the hybrid replicons into keratinocytes with the HPV18 genome, passing the cells five times and analyzing DNA extracted from each pass by Southern blot (Figure 5B). Both the HPV16/18 and HPV31/18 hybrid replicons replicated at low levels for the first two passes, and only the HPV31 replicon was maintained over long-term cell division. However, the copy number of the HPV31/18 replicon was strikingly low compared to the HPV18 replicon and could only be observed at later passages after extended exposure of the blots. qPCR measurement of the HPV18 genome and HPV18 replicon copy numbers showed that the HPV18 genome copy number was relatively consistent across all co-transfections. In contrast, the HPV16 and HPV31 hybrid replicons were severely reduced compared to that of HPV18; the copy number of the HPV31 hybrid replicon was barely detected above non-maintained replicon controls (Figure 5C).

**Figure 5.**
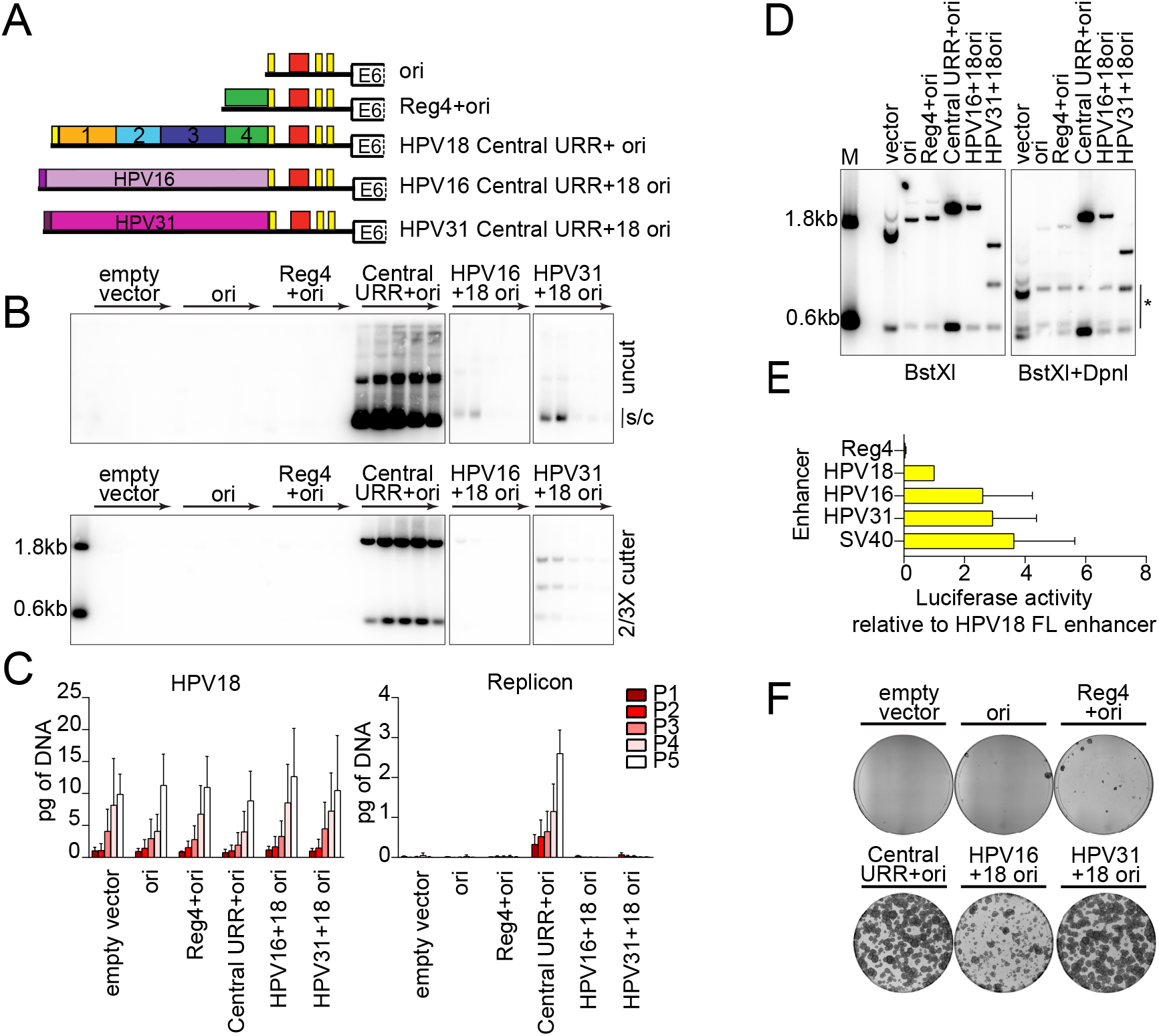
The central URR from HPV16 and HPV31 does not fully support HPV18 replicon maintenance. A. Diagram of enhancer replicons. B. DNA collected from five passages of co-transfected cells were analyzed by Southern blot as described in Figure 2C. The monomeric, supercoiled form (s/c) of the replicon is indicated. C. HPV18 genome (left) and replicon (right) copy numbers were measured by qPCR. D. DNA was collected from keratinocytes co-transfected with HPV18 and the indicated test replicon 48 hours after transfection. DNA samples were digested with *BstX*l, or *BstX*l and *Dpn*l, and visualized by Southern blot analysis as mentioned in Figure 2C. *Dpn*l digestion products are indicated by an asterisk. E. Keratinocytes were transfected with luciferase reporter plasmids containing the cis elements indicated as well as a positive control (SV40 enhancer) and negative control (empty luciferase vector containing a minimal promoter). Cell lysates were collected 48 hours after transfection and luciferase activity was measured and normalized to the total protein concentrations for each cell lysate and the negative control plasmid. Reporter activity for each enhancer is shown relative to the HPV18 full-length enhancer. The luciferase assays shown were conducted at the same time as those reported in Figure 4C and the activity of the positive control and HPV18 enhancer luciferase results are replicated here. F. Keratinocyte colonies arising from continuous G418 selection were stained with methylene blue approximately 14 days post-transfection. Error bars represent standard deviation. Data shown is representative (B, D, F) or an average of two (C) biological replicates. The average of three biological replicates is shown in E.

The low HPV16 and HPV31 test replicon copy number observed even at pass one suggested deficiencies in short-term replication. This was assessed by extracting DNA from cells at 48 hours posttransfection. As seen in Figure 5D, transient replication of HPV16 and HPV31 hybrid replicons was reduced compared to the HPV18 Central URR+ori replicon control.

Some studies have shown that the HPV18 URR has stronger enhancer activity compared to that of HPV16 and HPV31 [21, 22], while others show differences in enhancer activity in different cell types [23]. Although we find that enhancer activity did not correlate with replicon maintenance, we wanted to ensure that low enhancer activity was not responsible for the low replicon maintenance of the HPV16 and 31 hybrid replicons. Therefore, we tested the relative enhancer activities in our primary keratinocytes by cloning the central URR regions from HPV16 (nt 7463-7857) and HPV31 (nt 7456-7867) into the luciferase vector with the minimal promoter. As shown in Figure 5E, the HPV18 central URR enhanced luciferase activity about 41-fold, while HPV16 and HPV31 enhanced 82-fold and 100-fold, respectively. Therefore, enhancer activity strength could not explain the failure of the HPV16 and HPV31 hybrid replicons to be maintained robustly.

Replicons that are maintained at very low copy number, or are less efficient at partitioning, may be difficult to detect by Southern blot and qPCR analysis. Therefore, the colony forming assay was used to select for this type of replicon (Figure 5F). Under G418 selection the HPV31 hybrid replicon supported robust neomycin resistant colonies indicating that it was most likely maintained as an extrachromosomal plasmid. Conversely, cells co-transfected with the HPV16 hybrid replicon formed aborted, collapsing colonies indicating the replicon was not maintained long-term. Together, these results indicate that the HPV31 central URR, like HPV18, contains the minimum cis elements needed for replicon maintenance, but that other cis elements may be necessary to bolster initial replication and/or increased partitioning efficiency.

Our finding that the HPV18 replicon was established and maintained much more robustly than those of HPV16 or HPV31 is highly consistent with a study from Lace et al. [24]. Lace and colleagues showed that HPV18 could form keratinocyte colonies containing replicating HPV18 genomes much more efficiently than HPV16 or HPV31. They also showed that initial viral transcription and replication was equivalent among the different viruses and that the increased ability of HPV18 to form keratinocyte colonies did not depend on the tissue source of the keratinocytes [24]. Here, we confirm and extend this observation by showing that this increased efficiency of HPV18 genome establishment and maintenance maps to the central URR region.

### Transcription from defined promoter elements in the HPV18 replicon is unnecessary for replicon maintenance

The HPV18 major early P105 promoter overlaps the replication origin and, by binding to E2BS, the viral E2 protein represses P105 transcription by competing for binding with the cellular TBP protein to the adjacent P105 TATA box [25]. Other studies have shown that MAR based plasmids are only maintained extrachromosomally if they are transcriptionally active [26, 27].The HPV18 replicons also contain an SV40 enhancer-promoter element that drives expression of the neomycin resistance gene. To determine whether either of these elements are important for replicon maintenance, mutations were introduced to hinder transcriptional initiation from the HPV18 P105 promoter (TATA box mutation) and/or the SV40 promoter (~400bp deletion of the SV40 enhancer-promoter element; SV40enh-pro Δ) (Figure 6A). Shown in Figure 6B, mutation of the HPV18 P105 TATA box in the replicon did not affect neomycin resistant colony formation of transfected keratinocytes. In contrast, keratinocytes co-transfected with either of the SV40enh-proΔ replicons formed only abortive colonies, because expression of the neomycin resistance gene is dependent on the SV40 enhancer-promoter element.

**Figure 6.**
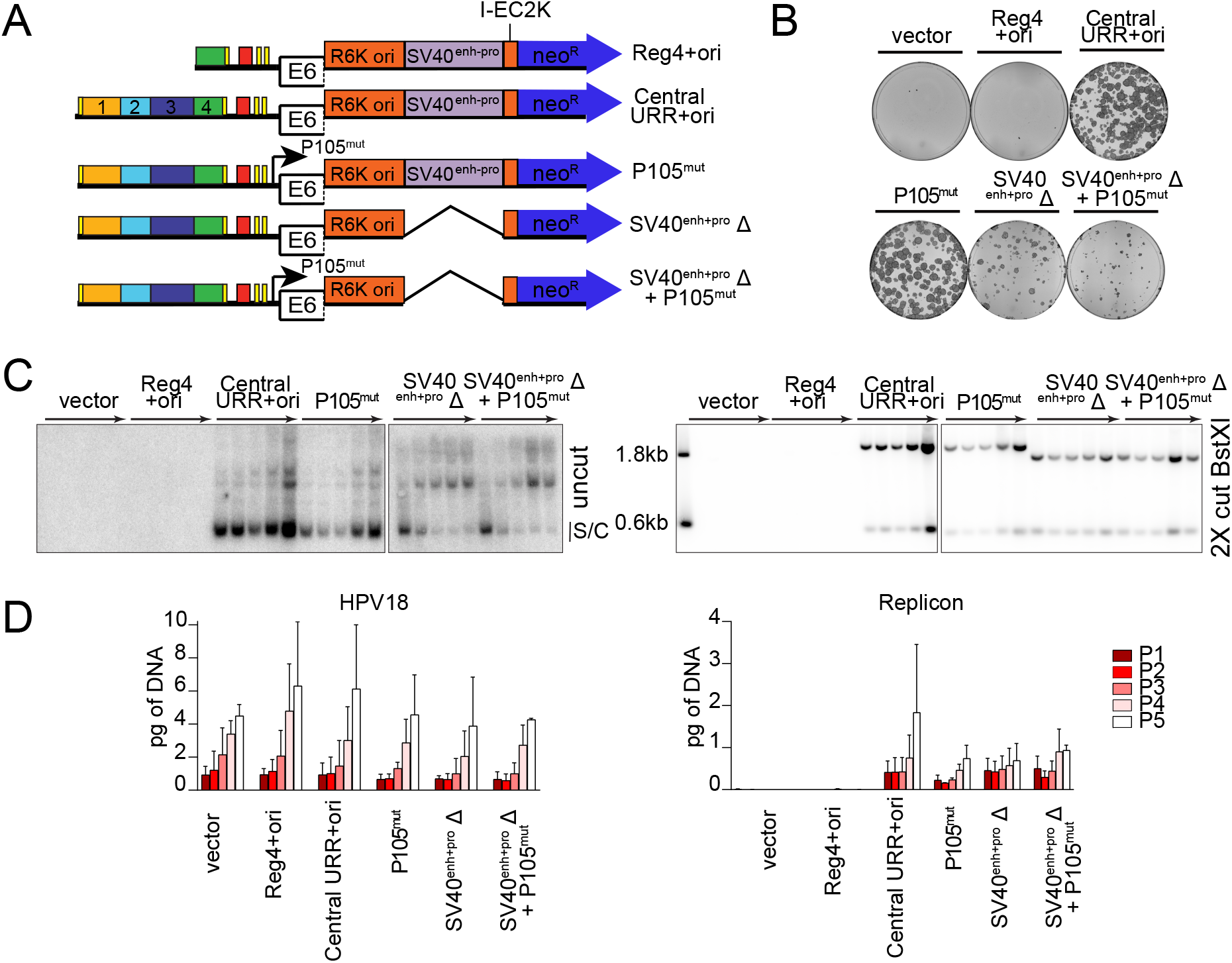
Transcription from the replicon is unnecessary for its extrachromosomal maintenance. A. Diagram of transcription initiation site mutations in replicons. B. Keratinocyte colonies arising from continuous G418 selection were stained with methylene blue approximately 14 days post-transfection. C. DNA collected from five passages of transfected cells were analyzed by Southern blot as described in Figure 2C. The monomeric, supercoiled form (s/c) of the replicon is indicated. D. HPV18 genome (left) and replicon (right) copy numbers were measured by qPCR. Error bars represent standard deviation. Data shown is representative (B, C) or an average (D) of two biological replicates.

To evaluate extrachromosomal maintenance of these replicons in the absence of G418 selection, DNA was collected from five passages of co-transfected keratinocytes and assessed by Southern blot analysis. As shown in Figure 6C, all three replicons were maintained over five cell passes. It was noted, however, that both SV40enh-pro Δ replicons formed higher molecular weight multimers as observed with the Reg2+4+ori replicons (Figure 3C). As proposed for the Reg2+4+ori replicons, the ~400bp deletion of the SV40 element may have negatively affected the topology or nucleosome spacing of the small circular DNA plasmid, or duplication of elements could result in more efficient replication or maintenance. Notably, RNA polymerase II recruits Topoisomerase I to relieve transcriptional-mediated torsional stress, and this could also promote replication (and perhaps resolution) of small circular molecules [28].

The copy number of the replicons and HPV18 genome was also assessed by qPCR (Figure 6D). The HPV18 genome copy number increased per pass and was comparable across all co-transfections. Similarly, the overall amount of all replicons increased per pass and was comparable to the pCGneo Central URR+ori positive control. Therefore, while the SV40 enhancer-promoter is important for expression of the neomycin resistance gene in the colony forming assay, neither this element nor the TATA box of the HPV18 P105 promoter was required for replicon maintenance.

### The 3’ half of region 2 modulates the replication efficiency/copy number of HPV18 replicons

To further define the function of Region 2 in replicon maintenance, a library of 13 linker scanning that spanned HPV18 nucleotides 7564-7641 was generated in the pCGneo Central URR+ori replicon, which lacks the CER element, a ColE1 bacterial element introduced to reduce replicon multimer formation (Figure 7A). Each linker replaced six bp of Region 2 with a Bgl*II* linker (AGATCT). Keratinocytes were cotransfected with the HPV18 genome and each of the 13 Bgl*II* replicons and were assayed in the colony formation assay with G418 selection (Figure 7B). All replicons gave rise to large numbers of neomycin-resistant colonies, indicating that no single linker mutation abrogated replicon maintenance. However, the centrally located Bgl linker replicon, Bgl 7, reproducibly resulted in fewer, less robust colonies, suggesting that this region might be important for establishment of the replicons.

**Figure 7.**
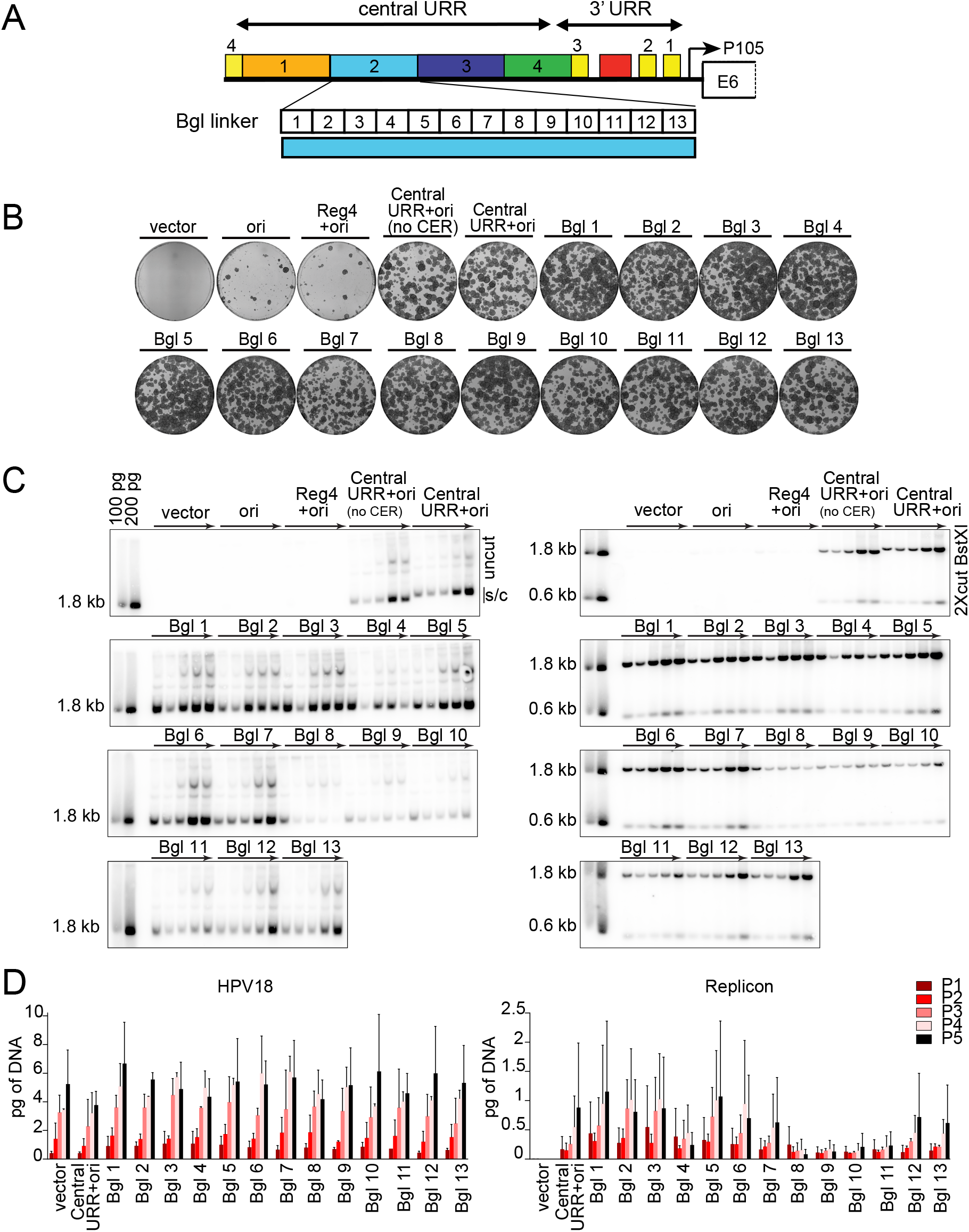
Linker scanning mutagenesis shows that the 3’ half of Region 2 modulates replicon copy number and/or partitioning efficiency. A. Diagram of the HPV18 central URR and ori. The location of Bgl linker scanning mutations introduced into HPV18 Region 2 are indicated by individually numbered white boxes (Bgl 1-13). All Bgl linker replicons lack the CER element (Figure 1B) and are compared to positive controls with and without a CER element. B. Keratinocyte colonies arising from continuous G418 selection were stained with methylene blue approximately 14 days post-transfection. C. DNA collected from five passages of transfected cells were analyzed by Southern blot as described in Figure 2C. The monomeric, supercoiled form (s/c) of the replicon is indicated. D. HPV18 genome (left) and replicon (right) copy numbers were measured by qPCR. Error bars represent standard deviation. Data shown is representative (B, C), or an average (D) of two biological replicates.

To confirm the extrachromosomal state of the Bgl linker replicons, DNA was extracted from keratinocytes passed five times after transfection without selection and analyzed by Southern blotting. As shown in Figure 7C, while all Bgl linker replicons were maintained extrachromosomally, replicons Bgl 8 through 11 were maintained at a reduced copy number. Measurement of the HPV18 genome and replicon copy number by qPCR confirmed that that the copy number was greatly reduced in Bgl 8 through 11 replicons and did not increase with pass (Figure 7D). Notably, the 3’ half of Region 2 is one of the most conserved regions among the central enhancer region of Alphapapillomavirus-7 HPVs (Figure 1C).

Together these results suggest that the 3’ half of region 2 contains elements that promote replicon establishment (Bgl 7 linker; nt 7600-7605) and long-term replication, copy number, or partitioning efficiency (Bgl 8-11 linkers; nt 7609-7629). No specific sequence in Region 2 seems essential for maintenance replication, but together sequences from the 3’ half of this region promote efficient establishment and longterm maintenance replication of the HPV18 replicons.

### No specific sequence in Region 2 is absolutely required for plasmid maintenance

To further analyze the requirement of specific sequences within Region 2, we used an “*in vivo* exonuclease assay”. Each of the 13 Bgl linker replicons, in addition to controls devoid of a *Bgl*ll restriction site (CER containing empty vector and wild-type HPV18 Central URR+ori replicon), were digested with *Bgl*ll and individually co-transfected with the HPV18 genome into primary keratinocytes. Within the cell the ends of the linearized replicon DNA are susceptible to cellular exonuclease activity before they are ligated by endogenous cellular enzymes. After transfection, the cells were passed four times without selection and low molecular weight DNA was isolated at the fourth pass. The recovered DNA was transformed into *E.coli* where the I-EC2K promoter drives expression of the neomycin resistance gene, which confers kanamycin resistance. Individual colonies were isolated and replicon plasmid DNA was extracted and sequenced. Replicons from a total of 864 bacterial colonies (approximately 66 colonies per bacterial transformation) were sequenced to determine the extent of any exonuclease induced deletions occurring at each *Bgl*ll site. In theory, Region 2 sequences that did not sustain any deletion were likely to be important for replicon maintenance in dividing cells. The recovered replicons contained 105 mutations (98 deletions and 7 insertions) and representative examples of these are shown in the sequence alignment in Figure 8A. Notably, however, no single sequence within Region 2 was protected from deletion in the recovered replicons. To determine whether deletions occurred more frequently in any part of the Region 2 element, the deletion frequency was calculated for each Region 2 nucleotide in the 98 recovered replicons that contained deletions (heatmap, Figure 8A). Nucleotides in the first half of Region 2 (overlapping Bgl linkers 1-7) were deleted in ~20-63% of the recovered replicons with deletions. In contrast, nucleotide deletions in the second half of Region 2 (overlapping Bgl linkers 8-13) only occurred in 1-14% of the recovered replicons. Thus, as shown for the BglI linker replicons in Figure 7, although no specific sequence element within Region 2 was absolutely necessary for replicon maintenance, fewer deletions are tolerated in the second half of Region 2 spanning Bgl linker 8-13.

**Figure 8.**
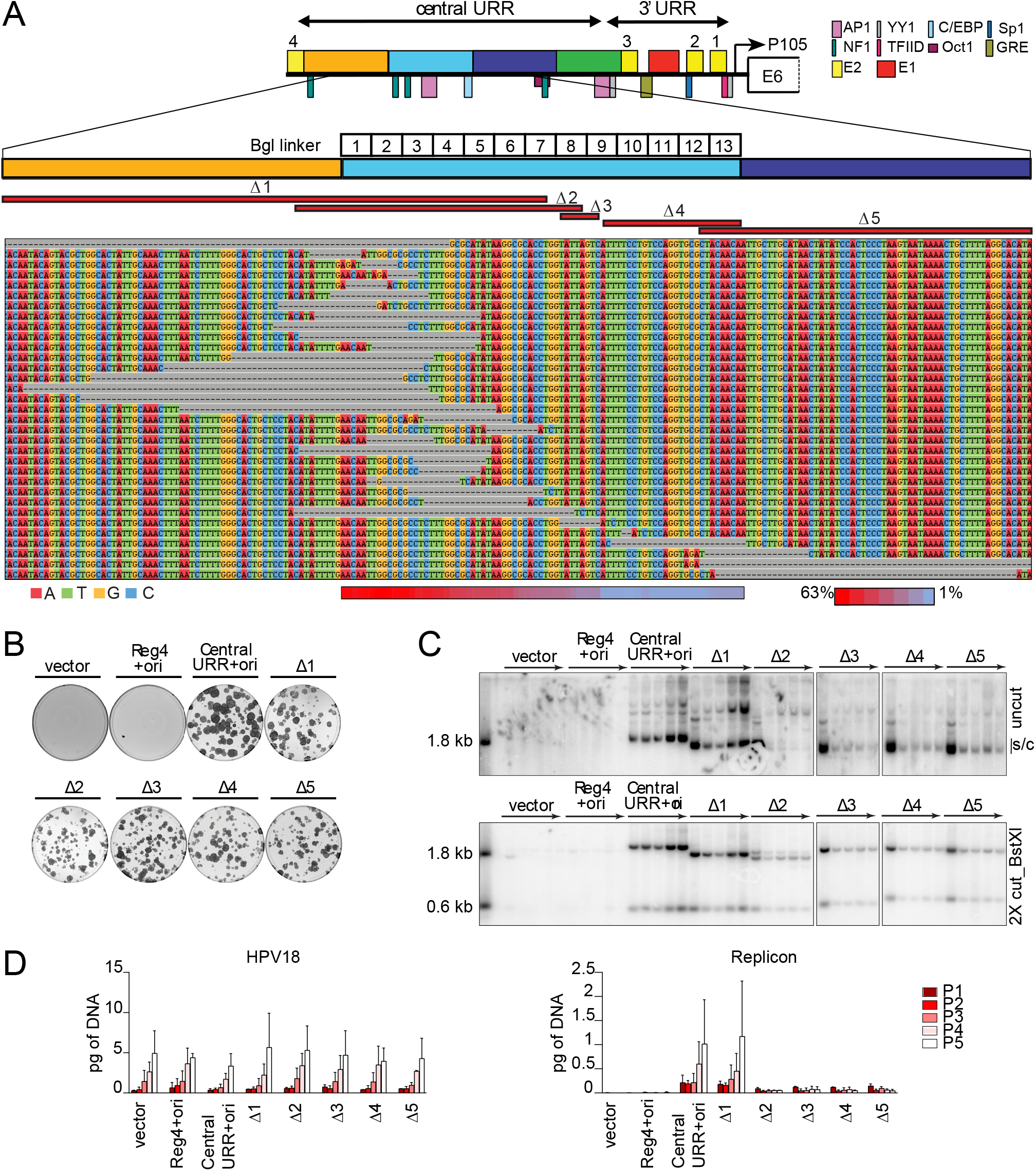
No single sequence in Region 2 is absolutely required for replicon maintenance but this region modulates replicon copy number and/or partitioning efficiency. A. Diagram of the HPV18 central URR and ori in addition to the location of important viral (E1, E2) and cellular transcription factor binding sites. The location of Bgl linker scanning mutations introduced into HPV18 Region 2 are indicated by individually numbered white boxes (Bgl 1-13). A representative sequence alignment of deletions present in Bgl linker replicons recovered from passaged cells. Deletions are highlighted by grey dashes while nucleotides are individually colored as indicated. Individual deletions present in select replicons recovered from the Bgl chewback assay are mapped by red bars (labeled Δ1 through 5) above the sequence alignment. The frequency with which a Region 2 nucleotide was deleted in the recovered replicons is indicated by a heatmap ranging from a maximum 63% (red) to 1% (blue) of the clones showing a deletion at a nucleotide site. B. Keratinocyte colonies arising from continuous G418 selection were stained with methylene blue approximately 14 days post-transfection. C. DNA collected from five passages of transfected cells were analyzed by Southern blot as described in Figure 2C. The monomeric, supercoiled form (s/c) of the replicon is indicated. D. HPV18 genome (left) and replicon (right) copy numbers were measured by qPCR. Error bars represent standard deviation. Data shown is representative (B, C), or an average (D) of two biological replicates.

To confirm that replicons isolated from the *in vivo* exonuclease assay could still replicate, five recovered replicons, collectively containing deletions across the HPV18 transcriptional enhancer (Δ1 - Δ5, mapped as red bars in Figure 8A), were tested for extrachromosomal maintenance in HFKs. Keratinocytes co-transfected with any of the Bgl Δ replicons formed neomycin resistant colonies, supporting results from the chew back analysis (Figure 8A), though they were not as robust or as numerous as the Central URR+ori control replicon (Figure 8B). In particular, although there were numerous colonies resulting from transfection of the Δ2 replicon, they were smaller in size, and many appeared abortive, compared to those resulting from transfection of the control replicon.

To ensure that the recovered Bgl Δ replicons were maintained extrachromosomally, they were cotransfected into keratinocytes with the HPV18 genome and cells were cultured for five passages without selection. DNA extracted from each pass was analyzed by Southern blot analysis. Although all Bgl Δ replicons were extrachromosomally maintained, the replicon copy number was severely reduced in keratinocytes transfected with Bgl Δ 2 – 5 and did not increase with pass like the central URR+ ori replicon (Figure 8C). Measurement of the HPV18 genome and replicon copy number by qPCR demonstrated that the HPV18 genome copy number was relatively uniform in all samples while the replicon copy number was greatly reduced in samples co-transfected with Bgl Δ 2 – 5 (Figure 8D). The reduced replicon copy number of Bgl Δ 2 – 5 replicons is similar to that observed with Bgl linker replicons 8-11 (Figure 7), which overlap in the same region. Also, of note is that the monomeric form of the Δ2 replicon decreased with pass until most of the replicon was present as a multimer, indicating further instability of this replicon. Together, these results imply that DNA elements within nucleotide 7606-7629 of Region 2 are important for *robust* replicon maintenance, but that no specific elements within Region 2 is absolutely necessary for maintenance replication.

### HPV18 ori proximal and distal AP1 binding sites promote both short-term and long-term replication

Little is known about the impact of specific transcription factors on HPV18 extrachromosomal genome maintenance. However, members of the AP1 family of transcription factors bind to the HPV18 central URR at conserved sites in Regions 2 (overlapping Bgl 8 and 9) and in Region 4, and this binding is critical for HPV18 gene expression [13]. To test the impact of AP1 on extrachromosomal maintenance, mutations previously shown to disrupt AP1 binding with minimal sequence change [13] were introduced into these sites in the Central URR+ori replicon (Figure 9A).

**Figure 9.**
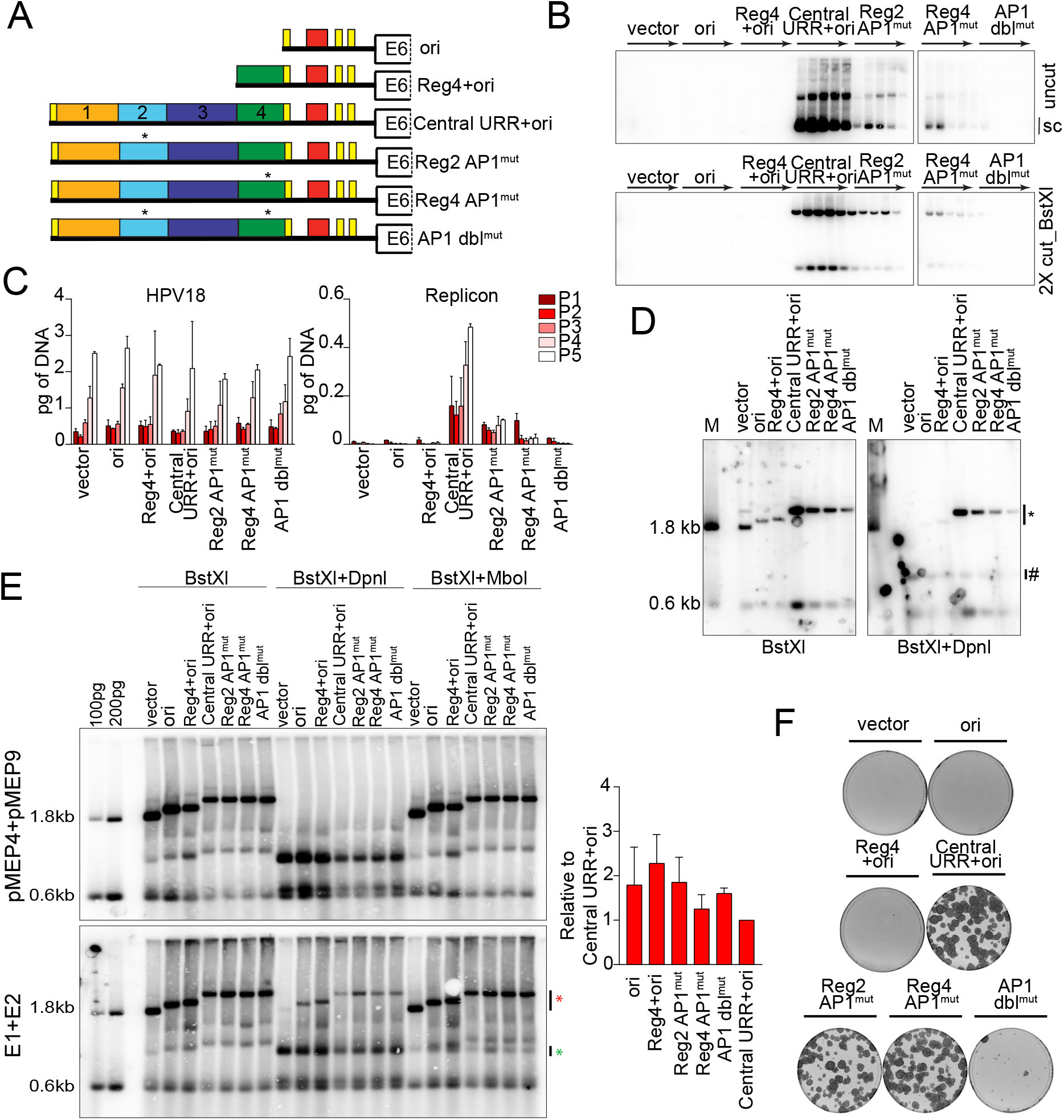
Mutation of AP1 binding sites impacts replicon replication. A. Diagram of AP1 mutations present in replicons. B. DNA collected from five passages of co-transfected cells were analyzed by Southern blot as described in Figure 2C. C. HPV18 genome (left) and replicon (right) copy numbers were measured by qPCR. D. DNA was collected from keratinocytes co-transfected with HPV18 and the indicated test replicon 48 hours after transfection, and analyzed by Southern blot as described in Figure 5D. Lanes 1-7 correspond to the following replicons: vector (1), HPV18 ori (2), HPV18 Reg4+ori (3), HPV18 Central URR+ori (4), Reg2 AP1^mut^ (5), Reg4 AP1^mut^ (6), and Reg2+4 AP1^mut^ (7). E. Keratinocytes were co-transfected with the indicated replicons and either empty pMEP vector control plasmids or pMEP E1 and E2 expression plasmids. E1 and E2 expression was induced with CdSO4 24 hours post-transfection, for five hours. Low-molecular-weight DNA was extracted 48 hours post-transfection and analyzed by Southern blot analysis as described in Figure 5D with an additional restriction digest that specifically digests replicating DNA (*Mbol*). *Mbol* sensitive and *Dpn*l resistant bands are indicated by green and red asterisks, respectively. *Dpn*l resistant DNA was quantified by phosphorimaging analysis of three independent experiments. Values shown in the graph are relative to the control Central URR+ori replicon. F. Keratinocyte colonies arising from continuous G418 selection were stained with methylene blue approximately 14 days post-transfection. Error bars represent standard deviation. Data shown is representative (B, E), or an average (C) of two biological replicates. Data shown for transient replication is representative of one (D) of three (E) bioreplicates.

The impact of the AP1 binding site mutations on replicon extrachromosomal maintenance was assessed by co-transfecting keratinocytes with HPV18 and replicon DNA, passing the cells five times, and analyzing DNA extracted at each pass by Southern blot. Each mutation severely hampered the levels of replication, but the DNA was maintained extrachromosomally at low levels for all five passes. When both sites were mutated (Reg2+4 AP1^mut^) the replicon DNA could not be detected after the first pass (Figure 9B). The copy number of the HPV18 genome and replicon were measured by qPCR (Figure 9C) and this confirmed that, while the HPV18 genome copy number was relatively consistent across all transfections, the replicon copy number was largely diminished in co-transfections with Reg2 or 4 AP1 mutations and barely detectable in Reg2+4 AP1^mut^ co-transfections. These results suggest at least one central URR AP1 binding site is necessary for replicon maintenance, but that AP1 binding at either site is sufficient for replicon replication.

The drastic drop in copy number suggested that the AP1 mutations might affect initiation of replication rather than replicon partitioning. Therefore, transient replication of the replicons was assessed 48 hours after co-electroporation of the replicons and HPV18 genome. Mutation of AP1 binding sites in either Region 2 or 4 greatly reduced short-term replication of the replicons, with this effect being further compounded in the Reg2+4 AP1^mut^ replicon (Figure 9D). This result implied that AP1 binding to the URR regulates initiation of HPV18 replication. To explore this further, the replicons were co-transfected with E1 and E2 expression plasmids (using liposome-mediated transfection). As shown in Figure 9E, in this case, all AP1^mut^ replicons replicated at levels comparable to the Central URR+ori control, as did the ori replicon. An explanation for these contrasting results is that high levels of E1 and E2 induce a DNA damage response [29, 30]; under these circumstances, viral DNA is transiently amplified but will not become established as the cells will not survive. Thus, all replicons can amplify transiently to equivalent levels in the presence of high levels of E1 and E2 proteins, however, only replicons complemented with the HPV18 genome can replicate long-term. This is most likely due to functions of the E6 and E7 proteins [31]. Therefore, the AP1^mut^ replicons show reduced copy number in long-term replication.

To further analyze the replication and maintenance of the AP1^mut^ replicons, keratinocytes were cotransfected with the HPV18 genome and replicon DNA and selected with G418 until neomycin resistant colonies formed. The individual Reg2 and Reg4 AP1^mut^ replicons supported robust neomycin resistant colonies that were comparable to the wild-type control (Figure 9F), however, keratinocytes co-transfected with the double AP1 site mutated Reg2+4 AP1^mut^ replicons failed to form colonies. Therefore, at least one AP1 binding site is necessary for establishment and maintenance of HPV18 replicons. However, due to the replication defects noted for Reg2+4 AP1^mut^ (Figure 9D), it is not clear whether loss of this replicon over long-term cell division is due to inhibited partitioning or dilution of the extrachromosomal replicon DNA over time.

## Discussion

We have previously shown that nucleotides 7564-7641 (Region 2) from the central URR, are essential *in cis* to the replication origin for maintenance of HPV18 derived replicons. In the current study we have further analyzed how these sequences might promote persistence of the replicons in dividing cells. We find that no single unique element or binding site is both necessary and sufficient for genome persistence, but several regions appear to be important for replicon copy number, establishment, stability, and the ability to persist as a monomer, as opposed to multimeric species. In agreement with our experimental findings, the most conserved part of Region 2 is nucleotides 7606-7641, which overlaps the Bgl8-13 linker mutations, seems to be most important for maintenance replication. For the most part, mutations in this region greatly reduce replicon copy number but individually do not completely eliminate long-term persistence of the replicon. Nucleotides 7604-7642 are 100% conserved among all 123 HPV18 variants sequenced to date (data not shown).

One transcription factor binding site that does seem to play an important, but indirect role on replication of the replicon is AP1. There is an AP1 binding site in the center of Region 2 (overlapping the Bgl8 and 9 linkers) and another in Region 4 and both sites are well conserved among all Alphapapillomavirus 7 HPVs. Both of these sites are important for viral transcription and AP1 has been proposed to be the “principal activator” of the HPV enhancers [16]. Notably, AP1 is a pioneer factor that can bind to and open repressed chromatin as well as activate transcription from HPV18 chromatin templates in vitro [32]. In this study, mutation of either AP1 site drastically reduced copy number of the replicon and we predict that these binding sites are structural opening/positioning elements that have long range effects on adjacent sequences. Notably AP1 enhances replication of SV40 and polyomavirus by interacting with TAg and enhancing unwinding of the origin [33]. Furthermore, AP1 (Jun-B/Fra-2) binds to the site in Region 2 of the HPV18 URR and cooperates, in specific helical phasing, with architectural protein HMG-I(Y) to form an enhancesome [34].

Many of the mutations generated in the URR (e.g. Bgl8-13 mutations) have dramatic effects on replicon copy number, but not on persistence. This copy number decrease was also observed previously when Region 3 (nt 7642-7749) was deleted. Because we were unable to identify a specific element or binding site important for replicon maintenance, we propose that HPV18 central URR elements, or the MMEE, is instead necessary for creating an open chromatin structure and controlling nucleosome positioning through the URR and into the origin. Nucleosome positions have been mapped in HPV16 and HPV18 [32, 35]: in both viruses a nucleosome is positioned in the enhancer region (the 5’ half of Region 2) and another over the origin of replication (there is about 90bp difference in the position on HPV16 vs HPV18). Nucleotide substitutions or deletions in these regions could have long range effects on adjacent sequences by disrupting this nucleosome positioning, resulting in chromatin-mediated repression.

Of note, nucleotides 7604-7642 contain an E-box (CAGGTG) that can be bound by MYC/MAX dimers [36] and is conserved among all Alphapapillomavirus 7 HPVs. C-myc is a universal amplifier of transcription [37] and promotes DNA replication [38]. While there is no direct evidence that C-myc binds to this motif in the HPV URRs, we speculate that it could be an additional way in which this region could promote robust replication and maintenance.

Homologous MMEE regions from the HPV16 and HPV31 URR regions could only partially complement replication of an HPV18 origin containing replicon. While it might be expected that these enhancer regions should have similar transcription factor binding sites, when considered in the context of the precise spacing of binding sites and nucleosomes, this observation is not unexpected. The overall organization of the HPV replication origin and upstream enhancer elements has been conserved among Alphapapillomaviruses but the precise location of transcription factor binding sites and position of nucleosomes has not [35]. There are cell-type specific differences in the transcriptional activity of different HPV URRs [23] and this could reflect subtle differences in tissue tropism between different groups of HPVs [39]. There are also large differences in the ability of different HPVs to replicate and establish their genomes as extrachromosomal plasmids in keratinocytes [24]. HPV18 genomes replicate and establish more efficiently than those of HPV16 and HPV31, although it is not understood why [40]. Our study suggests that these differences could be due to differences in the viral MMEE elements, rather than in expression or function of the viral gene products.

In our studies, some replicons persist, but the monomeric form diminishes per cell pass and multimeric forms become enriched, indicating that the monomeric form is not optimal. This could be due to restraints on topology, steric hinderance between binding factors when two elements are juxtaposed, or sub-optimal nucleosome positioning. Cellular transcription factors can promote loop formation between the HPV URR and other regions of the genome [41], and the E2 protein both bends DNA at the binding motif [42], and forms loops in the viral DNA by interacting with other bound E2 proteins [43]. Taken together, these long-distance interactions could greatly modulate the efficiency of DNA synthesis and partitioning of replicons.

It is generally accepted that HPV genome partitioning is mediated by the E2 protein, which tethers the viral DNA to host chromosomes to mediate viral genome retention and partitioning [3]. However, the paucity of E2BSs in the Alphapapillomavirus HPVs, and the finding that only those E2BSs in the replication origin are important for these processes [8, 9], initiated a search for cellular factors that could mediate or strengthen tethering to host chromosomes by binding to motifs in the MMEE. Based on the findings of this current study however, we propose that the role of the MMEE is to strengthen protein-protein interactions at the replication origin by maintaining an active and open chromatin environment. The cellular bromodomain-containing protein 4 (Brd4) binds to the E2 protein and regulates papillomavirus transcription [44, 45] and this is mediated by the E2BS in the replication origin [32]. Brd4 co-localizes with the E2 protein from many papillomaviruses on mitotic chromosomes [46–48]. However, the interaction of Brd4 with Alphapapillomavirus E2 proteins is weak and difficult to observe on mitotic chromosomes [49] and mutation of key E2 residues necessary for Brd4 interaction does not inhibit maintenance of Alphapapillomavirus genomes [50]. Nevertheless, the low affinity interaction of Alphapapillomavirus E2 proteins and Brd4 observed in vitro, or when expressed in the absence of viral DNA, could be stabilized by the interactions described here. Notably, Delcuratolo and colleagues have shown that Brd4, in complex with the SfPV1 (cotton tail rabbit papillomavirus) E2 protein activates expression of the AP1 family member c-fos [51]. In turn, c-fos promotes viral transcription and, we predict based on the results of our study, augments viral replication. Additionally, we have previously shown that Brd4-E2 complexes associate with active regions of host chromatin [52]. Hence, elements in the HPV MMEE element that promote active chromatin configurations could further enhance these interactions with host chromosomes.

In summary, we have used highly robust, but sensitive, complementation assays to determine the requirements for HPV18 derived replicon maintenance. These assays can both detect very low copy number replicons (colony formation assay) and monitor the presence of monomeric, supercoiled molecules over many keratinocyte population doublings (Southern blot analysis). We conclude that the main function of the previously defined MMEE element is to enhance and promote an active chromatin configuration that is conducive to long-term replication and partitioning.

## Materials and Methods

### Alignment of Alphapapillomavirus 7 central URR sequences

URR sequences from the nine reported Alphapapillomavirus 7 species were retrieved from the PaVE website (https://pave.niaid.nih.gov) [53] and aligned in Geneious Prime using the multiple sequence Geneious global alignment algorithm with free end gaps (65% similarity cost matrix [5, –4]). Mean pairwise identity scores shown are were calculated using a sliding window size of 1.

#### Plasmids

HPV18 replicons were generated in the pCGneo plasmid described previously [8]. Elements were inserted between the CER element and the HPV18 ori or Reg4 element using standard cloning procedures. Despite having a CER element to prevent the formation of multimers in bacteria, some replicons had the tendency to multimerize. Therefore, monomeric supercoiled plasmids were always gel purified before transfection. pMEP4-HPV18 E2 and pMEP9-HPV18 E1 plasmids were described previously [54].

A series of 13 Bg*l*ll linker mutations were introduced between HPV18 nucleotide 7564 to 7641 in pCpG-URR 7452-174 replicon using PCR. This replaced every six nucleotides with the sequence AGATCT. Bgl Δ replicons were rescued from cells transfected with the Bgl*ll* linker plasmids, as described in the results. These contained deletions in pCpG-URR 7452-174: Δ1 = nt 7455-7602, Δ2 = nt 7555-7608, Δ3 = nt 7606-7613, Δ4 = nt 7613-7638, Δ5 = nt 7633-7749).

Enhancer regions were cloned upstream from the HPV18 origin in pCGneo 7752-174. The K5 enhancer was amplified from cellular DNA and contains sequences from chr12 52522417-52522989. The 279bp human AldA enhancer was PCR amplified from the pDRIVE5-GFP-10 plasmid (Invivogen). The HCMV enhancer was PCR amplified from the pDrive5 GFP-1 Promoter Test (Invivogen). The E2BS4 and enhancer regions from HPV16 (nt 7451-7857) and HPV31 (nt 7477-7867) were amplified by PCR and cloned into pCpG-URR 7817-174 at the indicated Nco*l* site in Figure 2A. The resulting plasmids are named hAldA+Reg4+ori, HCMV+Reg4+ori, K5+Reg4+ori, HPV16 Central URR+18ori, and HPV31 Central URR+18ori, respectively.

All enhancer elements were amplified by PCR and inserted into the Xho*l* and Hind*lll* sites of the Promega pGL4.23 vector. This vector expresses luciferase from a minimal promoter consisting of a TATA element and a transcriptional start site. The resulting plasmids are named pGL4.23 HCMV, pGL4.23 hAldA, pGL4.23 K5-F, pGL4.23 K5-R, pGL4.23-HPV16 Central URR, pGL4.23-HPV31 Central URR, pGL4.23-HPV18 Reg4, pGL4.23-HPV18 Reg2+4, and pGL4.23-HPV18 Central URR+ori. The pGL4.13 vector that expresses luciferase from the SV40 enhancer/promoter was used as a positive control.

In indicated replicons, the HPV18 P105 TATA box was mutated using site directed mutagenesis at HPV18 nt 76-79, converting TATATAAAA to TATATttAt. The SV40 enhancer-promoter element was deleted from select HPV18 replicons using *Sbfl* and Hind*lll* restriction digest and blunt ended ligation. Lastly, previously characterized HPV18 central URR AP1 binding site mutations [55] were introduced using PCR amplification and standard restriction digest cloning. These minimal mutations abrogate AP1 binding without affecting adjacent factors.

All oligonucleotide sequences are in Supplementary Table 1.

#### Cell culture

Primary human keratinocytes were co-cultured in Rheinwald-Green F medium (3:1 Ham’s F-12/high-glucose Dulbecco’s modified Eagle’s medium [DMEM], 5% fetal bovine serum [FBS], 0.4 μg/ml hydrocortisone, 8.4 ng/ml cholera toxin, 10 ng/ml epidermal growth factor, 24 μg/ml adenine, 6 μg/ml insulin) with lethally irradiated J2-3T3 murine fibroblasts. HFKs were maintained in 10μM Y-27632 until ~24 hours post-transfection. For experiments using G418 selection, HFKs were cultured on G418-resistant J2-3T3 murine fibroblasts.

#### Transfection

Electroporation was performed using the Amaxa electroporation system (Lonza) according to the manufacturer’s instructions. Briefly, 1X10^6^ cells were mixed with 2 μg of DNA (1 μg of HPV18 genome and 1 μg of the HPV18 replicon) and electroporated using the T-007 program (optimal for enhanced keratinocyte survival). A total of 1.4X10^6^ electroporated cells were plated on irradiated feeders for transient replication experiments, 0.4X10^6^ cells were plated for long-term replication experiments, and 0.1X10^5^ were plated for colony formation experiments. The latter group of cells was plated on irradiated G418-resistant J2-3T3 cells, selected in 400 μg/ml G418 for 4 days, and then in 200 μg/ml G418 until colonies appeared. The cells were fixed and stained with methylene blue (14-18 days post transfection).

#### E1 and E2 transient replicon replication

Six well plates containing ~2.5×105 cells per well were transfected using Lipofectamine 3000 per the manufacturer’s instructions. Cell were co-transfected with a tested replicon and empty control plasmid pMEP4 and pMEP9 or pMEP9-E1 and pMEP4-E2 eukaryotic expression plasmids at molar ratio of ~4.5-4.9. E1 and E2 expression was induced ~24 hours posttransfection with 3uM of CdSO4 for five hours. Cell pellets were collected 48 hours post-transfection and low-molecular-weight DNA was extracted for Southern blot analysis.

#### DNA extraction

Whole-cell DNA extraction was carried out using the Qiagen blood and tissue kit according to the manufacturer’s instructions. Low-molecular-weight DNA was extracted using a modified Hirt extraction method [5].

#### Southern Blotting

Total cellular DNA was digested with either a restriction enzyme to cleave the viral DNA and derived replicons (*BstX*I; 2-3 cuts depending on the replicon) or a non-cutter (*Bgl*ll or *Nde*l) to linearize only cellular DNA. For transient DNA replication analysis, samples were also digested with *Dpn*l to identify *Dpnl*-resistant replicated DNA or *MboI* to specifically digest replicating DNA. After digestion, samples were separated on Tris-acetate-EDTA (TAE)-agarose gels. DNA was visualized with 10 μg/ml ethidium bromide and transferred to Nytran SPC membranes with a TurboBlotter downward transfer system (Whatman). Membranes were UV cross-linked, dried, incubated with prehybridization blocking buffer, and then incubated overnight with 25 ng [^32^P]dCTP-labeled HPV18 DNA or pCpG vector in hybridization buffer (3× SSC [1× SSC is 0.15 M NaCl plus 0.015 M sodium citrate], 2% SDS, 5× Denhardt’s solution, 0.2 mg/ml sonicated salmon sperm DNA). Radiolabeled probe was generated by purifying a digested (*Nco*l; 1 cut) pCpG vector DNA using High Pure PCR product Purification Kit (Roche) followed by radiolabeling the probe using Random Prime DNA labeling kit (Roche). Hybridized DNA was visualized and quantified using a Typhoon scanner (GE Bioscience).

#### Quantitative PCR for copy number assessment

Fifteen nanograms of whole-cell DNA was analyzed by qPCR using 300 nM primers and SYBR green master mix (Roche). The reaction conditions consisted of a 15-min 95°C activation cycle, 40 cycles of 10 s at 95°C for denaturation and 30 s at 60°C for annealing, and elongation. Copy number was calculated by comparison to standard curves of linearized HPV18 or pCGneo vector DNA. The RPPH1 DNA copy number was used as a normalization control. The primer sequences used are listed in Supplementary Table 1.

#### Luciferase assay

12 well plates containing ~ 0.13X10^6^ cells/well were transfected using Lipofectamine 3000 per the manufacturer’s instructions. Cell lysates were collected 24 hours post-transfection using Passive lysis buffer (Promega). Samples from three technical replicate transfections were measured in duplicate using a Zylux FB12 luminometer and the Dual-Luciferase Reporter Assay System (Promega). Luciferase activity measurements were normalized to total protein concentrations detected in cell lysates. Protein concentrations were determined using BCA assay (Pierce). Background levels of luciferase activity present in the negative control (empty luciferase plasmid with a minimal promoter) were deducted from the normalized measurements. Shown luciferase activity measurements are relative to the full-length HPV18 enhancer.

#### In vivo exonuclease assay

Bgl linker clones 1 – 13 and control plasmids (pCGneo vector and pCGneo-7452-174) were digested with *Bgl*ll and the ends dephosphorylated with Antarctic phosphatase. HFKs were electroporated with 1μg HPV18 DNA and 1μg digested pCGneo replicon and cells were cultured for four passes in the absence of G418 selection. Low molecular weight DNA was isolated from the third pass of transfected cells using a modified HIRT procedure [5], transformed into E. coli GT115 (Invivogen) with kanamycin selection. The plasmids recovered from approximately 66 bacterial colonies from each Bgl linker transformation were sequenced. Sequences were aligned with DNASTAR Lasergene Seqman Pro.

For the heatmap, the nucleotide deletion frequency for Region 2 was calculated by dividing the total number of times a nucleotide was deleted by the total number of replicons containing a deletion (98). Conditional formatting in excel was used to generate a heatmap that ranged between a maximum (63%) and minimum (1%) frequency. Maximum and minimum values are marked in red and blue, respectively.

### Ethics Statement

Primary human keratinocytes were isolated from anonymized neonatal foreskins provided to the Dermatology Branch at NIH from local hospitals. The NIH Institutional Review Board (IRB) approved this process and issued an NIH Institutional Review Board waiver.

## Acknowledgements

We thank all members of the McBride laboratory for helpful discussions. This work was supported by the Intramural Research Program of the National Institute of Allergy and Infectious Diseases at the National Institutes of Health.

